# Serological screening in animals combined with environmental surveys provides definite proof of the local establishment of *Burkholderia pseudomallei* in Guadeloupe

**DOI:** 10.1101/2024.02.09.579440

**Authors:** Mégane Gasqué, Vanina Guernier-Cambert, Gil Manuel, Rachid Aaziz, Jules Terret, Thomas Deshayes, Xavier Baudrimont, Sébastien Breurec, Emma Rochelle-Newall, Karine Laroucau

## Abstract

**Background:** Melioidosis is an emerging infectious disease caused by the soil-dwelling bacterium *Burkholderia pseudomallei* that affects both humans and animals. It is endemic in South and Southeast Asia, and northern Australia, causing an estimated 165,000 human cases annually worldwide. Human cases have been reported in the French West Indies (Martinique and Guadeloupe) since the 1990s. Conversely, no human cases have been reported in French Guiana, a French territory in South America. Our study aimed to investigate whether *B. pseudomallei* is locally established in Guadeloupe and French Guiana. We assessed animal exposure by serology and examined the presence of *B. pseudomallei* in the environment of seropositive animals.

**Methodology/Principal findings:** Blood samples were collected from domestic animals in two goat farms in Les Saintes, Guadeloupe (n=31), and in 56 farms in French Guiana (n=670) and tested by ELISA. Serological follow up was performed on selected farms. Soil, water and goat rectal swabs were collected and analysed by culture and PCR. In French Guiana, the highest prevalence rates were observed in equids (24%) and cattle (16%), while in Les Saintes, a prevalence of 39% was observed in goats. The longitudinal study in Les Saintes revealed consistent high seropositivity in goats. A *B. pseudomallei* strain was isolated from the soil from one of the farms and detected in goat rectal swabs from the other farm.

**Conclusions/Significance:** Our environmental investigation prompted by the serologic data confirms the presence of *B. pseudomallei* in Les Saintes, consistent with documented human cases of melioidosis on this island. In French Guiana, our serologic results call for environmental surveys and a re-evaluation of human infections with melioidosis-like symptoms. The approach developed in this study may help to identify high-risk areas that warrant further investigation.

**Author summary:** Burkholderia *pseudomallei*, an environmental bacterium, is the causative agent of melioidosis in humans and animals. If the disease has been historically reported to be endemic in South Asia and northern Australia, recent studies reveal its presence outside of these territories, both in the environment and among patients who have not travelled to endemic areas. Furthermore, the projected increase in extreme climatic events in the near future could increase the prevalence of the disease as well as cause its emergence in new territories. For these reasons, it is important to identify new areas at risk.

Our study aimed to investigate the presence of the pathogen in French West Indies. We combined surveys in domestic animals (cattle, goats, horses, sheep, and pigs) and in the environment. The identification of seropositive animals without clinical signs, together with the isolation of *B. pseudomallei* in the environment of a goat farm in Guadeloupe, underscores the importance of including melioidosis in animal surveillance programs. The use of serologic methods can help identify animal exposure to the pathogen, thereby helping to identify areas where the pathogen may be present in the environment.

---

**Keywords:** serology, veterinary science, ELISA, environment, melioidosis, *Burkholderia pseudomallei*

## Introduction

Melioidosis is an opportunistic infectious disease with a significant prevalence in South Asia, Southeast Asia, and northern Australia, primarily due to environmental exposure to *Burkholderia (B.) pseudomallei*. This telluric bacterium can affect both humans and animals through inhalation, ingestion, or skin contact [1]. It is listed as a Class 1 pathogen by the U.S. Centers for Disease Control and Prevention (CDC) and as a selected agent under the Microorganisms and Toxins (MOT) regulations by the French National Agency for the Safety of Medicines and Health Products (ANSM).

The global melioidosis burden was predicted to be 165,000 human cases in 2015 (2). Moreover, reports of human and, occasionally, animal cases are increasing along with environmental evidence of the bacteria worldwide, highlighting the probable presence of the bacterium in all continents, including Africa and the Americas [3–5].

The clinical presentation of the disease is highly variable between individuals and species, ranging from chronic forms to acute septicaemia [1,6]. Diagnosis and treatment of human melioidosis are complicated by the lack of pathognomonic clinical features and a long treatment duration. Several risk factors, such as diabetes, and chronic diseases, contribute to increased susceptibility [7]. In animals, melioidosis affects a variety of species with multiple cases documented in domestic animals such as cats, cattle, deer, dogs, goats, equids, sheep, and sporadic cases in other species, including marine mammals, birds and even crocodiles in zoos or wildlife parks [6].

Human melioidosis is a neglected disease and, as such, lacks accurate epidemiological data and diagnostic tools. These are even more limited for animal melioidosis, and the diagnostic approaches for animal melioidosis are generally the same as those in humans [6,8,9]. Although diagnostic methods such as indirect hemagglutination (IHA), complement fixation test (CFT) and enzyme-linked immunosorbent assay (ELISA) have been developed for commonly tested species such as small ruminants, pigs, and equids [10–12], standardised tests for the diagnosis of melioidosis in all susceptible species are currently lacking. A recent study highlighted the effectiveness of serological tests developed for glanders (a disease caused by *B. mallei*, a bacterium closely related to *B. pseudomallei*) for accurately detecting melioidosis in equids. These tests include both a reference method (CFT) and alternative methods (ELISA, Western blot) [13].

The bacterium *B. pseudomallei* is primarily found in moist clay soils and in turbid waters [14,15]. Its distribution in soils is strongly influenced by climatic events, with more cases observed during the rainy season and sporadic contamination peaking during extreme weather events such as storms or floods [16,17]. Environmental detection of *B. pseudomallei* is challenging and specific culture protocols are required to isolate the bacterium [18]. Its detection could be harder in non-endemic areas where the bacterium may be present at lower levels. A new culture medium, with erythritol as a carbon source, was recently developed to improve culture isolation from soil samples [19]. It was successfully used to isolate *B. pseudomallei* from a rice farm in South-Central Ghana, Western Africa [20].

Cases of melioidosis have been reported in both South, Central and North America. This includes the French West Indies, i.e., Guadeloupe and Martinique, where 21 human cases have been recorded since 1993 [21–25], some of whom had no history of travel to endemic areas. In contrast, no human cases have been reported in French Guiana (Guiana Shield, South America), while human cases have been reported in neighboring Brazil [3], as well as in Venezuela and Colombia [26].

The objective of this study was to demonstrate the local establishment of *B. pseudomallei* in French Guiana and Guadeloupe. We aimed to document animal exposure through a serology study, to document their immune response over time, and to investigate the link between seropositive animals and the presence of *B. pseudomallei* in their close environment. The serological analysis was performed with an ELISA kit – using a recombinant protein and double antigen technology – originally developed for the diagnosis of equine glanders [11] and was used in our study on different domestic species (goats, cattle, equids and sheep).

## Materials and Methods

### Animal sampling in French Guiana and Guadeloupe

In French Guiana, a total of 56 farms were surveyed, primarily along the northern coast where 80% of the population resides (Fig. 1A). These farms included both single-species and multispecies farms, and 2 to 31 animals per farm were analysed. Among the different regions, 24 farms were located in Macouria, 14 in Kourou, 9 in Mana, 7 in Saint Laurent du Maroni, 5 in Sinnamary, 3 in Montsinery Tonnegrande, 2 in Saint George de l’Oyapack and Cayenne, and 1 in Roura and Iracoubo. Notably, 10 of these farms had multiple species, including cattle, sheep and equids. In Les Saintes, part of the Guadeloupe archipelago, two goat farms were sampled, Farm A with approximately 40 goats and Farm B with approximately 60 goats on the island of Terre de Haut (Fig. 1B). On both farms, mixed-sex goats grazed freely in open pastures, consuming local vegetation, and no birth control measures were used, hence the approximate herd sizes reported by the farmers. The topography of the pastures was characterized by steep terrain and both were susceptible to significant surface water runoff.

**Fig 1.**
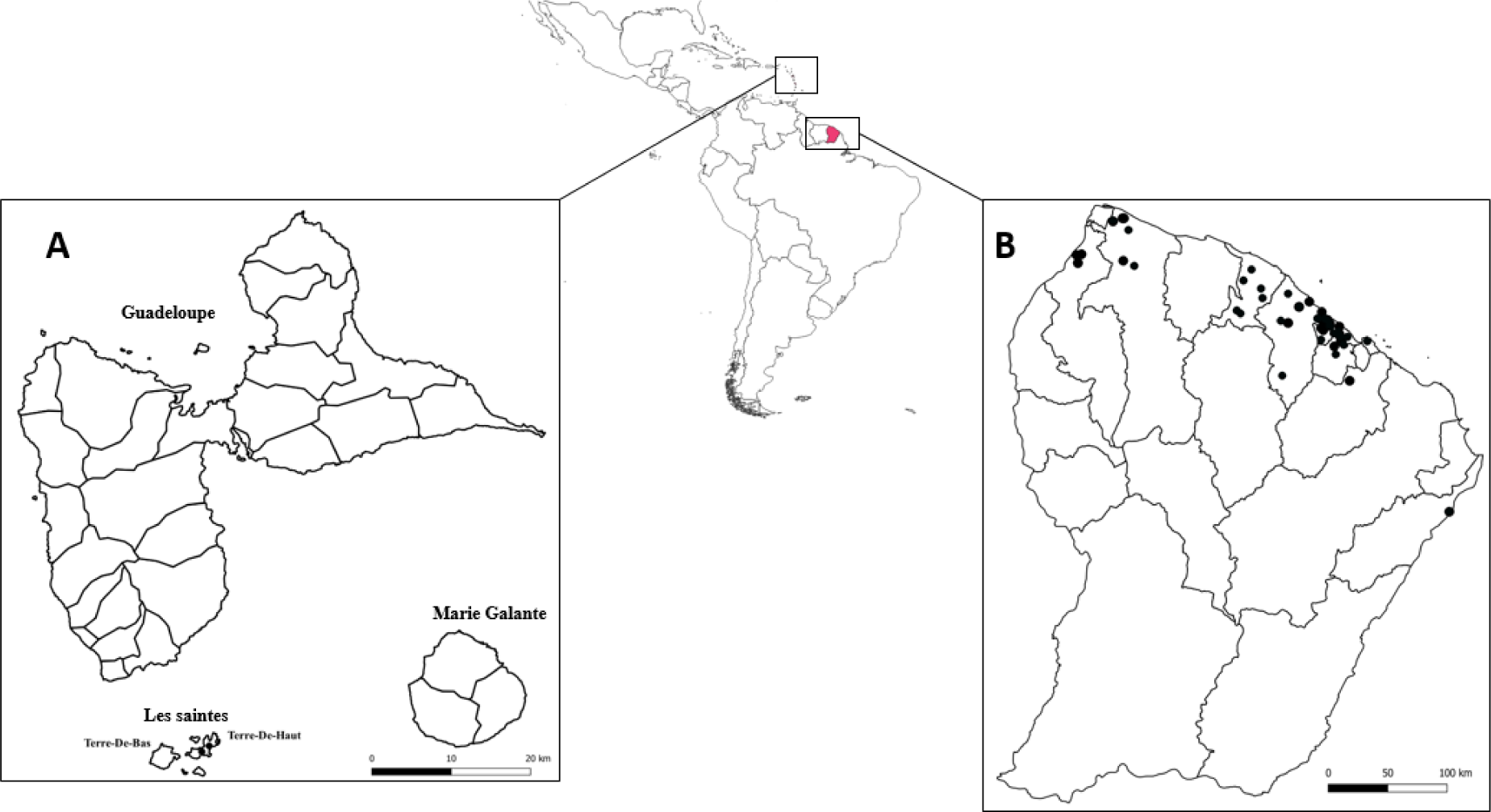
Map of sampling sites. Farms where animals were tested in (A) Guadeloupe and (B) French Guiana are represented as black dots.

Blood samples were collected between November 2021 and December 2022 from i) cattle (361 animals from 37 farms), sheep (131 animals from 15 farms), goats (100 animals from 10 farms), equids (63 animals from six farms), and pigs (15 animals from one farm) in French Guiana, and ii) goats (31 animals from two farms) in Les Saintes.

After blood clotting and centrifugation at 900 rpm for 10 min, sera were collected, heat-inactivated at 60°C for 30 min and stored at −20°C until analysis.

In some farms with GLANDA-ELISA positive animals (see § Serological analysis hereafter), serological longitudinal monitoring was conducted. In French Guiana, additional serological tests were conducted in one equid farm between October 2021 and January 2022. In Les Saintes, a more detailed longitudinal study was conducted in the two goat farms. Blood samples were collected between November 2021 and March 2023 (farm A) or December 2022 (farm B) (Fig 2). Rectal swabs were collected in duplicate for each animal between April 2022 and December 2022 (Fig 2). One swab was stored dry at −20°C until PCR analysis, while the second was stored in Luria-Bertani (LB)−20% glycerol at −20°C for culture.

**Fig 2.**
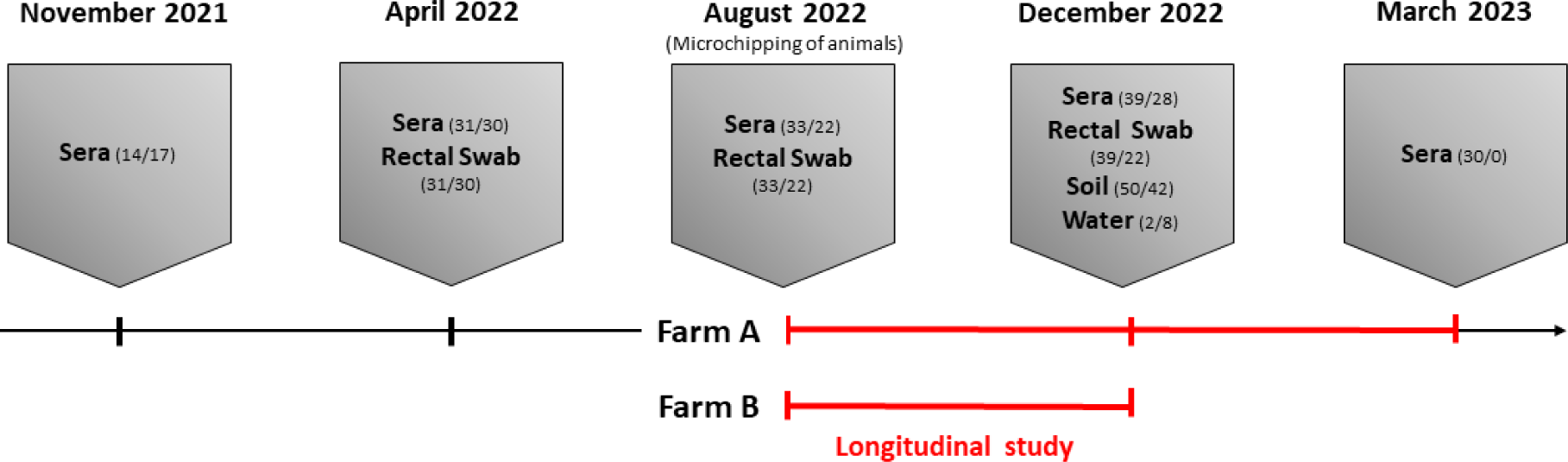
Sampling timeline in two goat farms in Les Saintes, Guadeloupe. Sampling occurred between November 2021 and March 2023. The longitudinal serological study (indicated in red) took place between August 2022 and March 2023 for farm A, and between August 2022 and December 2022 for farm B. The numbers in parentheses correspond to the number of samples collected from farms A and B, respectively.

Animal sampling protocols were approved by the IRD Ethics Committee (21 April 2022).

### Environmental sampling in Guadeloupe

Soil and water samples were collected from both goat pastures (farms A and B) in December 2022 during the rainy season (Fig 2). Soil samples (n = 50 in farm A, n = 42 in farm B) were collected every 5 m along a series of transects perpendicular to the main slope of the pasture. Each transect was distant by at least 6 m from the previous transect. Samples were collected 30 cm deep with a standard soil auger that was disinfected with absolute ethanol after each sampling. Approximately 200 g of soil was collected and placed in closed Ziploc bags to maintain moisture. Samples were stored at room temperature (approximately 25°C) and protected from light until shipped to France. Samples were analysed within two weeks of collection (see § Environmental sample processing). Water samples were collected in farm A (n = 2) and B (n = 8) in goat drinking troughs.

All samples (both animal and environmental) were sent to Anses, Maisons-Alfort, France, to be processed.

### Serological analysis

The ID Screen^Ⓡ^Glanders Double Antigen Multi-species ELISA test (Innovative Diagnostics, Grabels, France) (hereafter referred to as GLANDA-ELISA) was used according to the manufacturer’s instructions. This test does not rely on the use of a specific conjugate and can therefore be used for multiple animal species. The relative amounts of antibodies in serum samples were calculated by reference to the positive and the negative controls provided in the kit. The serological titer (expressed as the Sample to Positive ratio or S/P%) was calculated using the optical density (OD) reading at 450 nm following the formula: S/P% = ((OD sample - OD negative control) / (OD positive control - OD negative control)) x 100. Serum samples with S/P% <70% were considered negative and ≥70% were considered positive.

### Additional serological tests on equine sera, French Guiana

The complement fixation test (CFT), the reference method for glanders in equids, was performed as previously described using the cold method with the Malleus CFT antigen (Bioveta, Czech Republic) [27]. Serum samples were initially tested at 1/5 dilution. Samples with 100% hemolysis were considered negative, those with 25–75% hemolysis were considered doubtful, and those with 100% inhibition of hemolysis were considered positive. All samples primarily identified as suspect or positive were retested over a range of dilutions. The “titer” is a six-digit barcode corresponding to the intensity of hemolysis inhibition (0 = 0%; 1 = 25%; 2 = 50%; 3 = 75%; 4 = 100%) at the reciprocal dilutions (1/5, 1/10, 1/20, 1/40, 1/80, 1/160) for each sample. Anti-complementary activity (due to incomplete elimination of complement proteins during the serum heat inactivation) was checked for each serum. Additionally, the Luminex assay, originally developed for the diagnosis of glanders in equids, was performed as previously described using the heat shock protein (GroEL) (BPSL2697) and hemolysin-coregulated protein (Hcp1) (BPSS1498) antigens [28]. MRI#1 was used as positive control to determine S/P%, which was calculated for each antigen using the same formula as in the GLANDA-ELISA. S/P% values greater than 45% for GroEL and 43% for Hcp1 were considered positive.

### Environmental sample processing

Each soil sample was homogenized, and 10 g of soil was diluted in 10 mL of sterile water. After vigorous vortexing, the suspension was shaken on an orbital shaker at 160 rpm for one hour and then allowed to settle for 10 min. From the supernatant, 1 mL was centrifuged at 8,000 rpm for 10 min and the resulting pellet was stored at −80°C until PCR analysis and 1 mL was used for a two-step enrichment. First, 1 mL of supernatant was mixed with 9 mL of liquid Ashdown medium [29] and incubated at 37°C for 48 h. Second, the Ashdown enrichment was mixed, and 1 mL was transferred to 9 mL of erythritol medium [19] and incubated at 37°C for 96 h. One mL of erythritol enrichment was centrifuged at 8,000 rpm for 10 min and the resulting pellet was stored at −80°C until PCR analysis. The remaining 9 mL of enriched medium was stored at 4°C.

Within two hours of collection, 50 mL of each water sample were filtered sequentially through a 3 µm filter followed by a 0.2 µm filter. The 0.2 µm filters were folded in four, preserved in Lysogeny Broth (LB) (10 g peptone, 5 g yeast extract, 10 g NaCl for 1 L)-glycerol 20% and stored at −80°C until analysis. For analysis, the LB-glycerol 20% was discarded and the filters were aseptically cut in half. One half was placed in 10 mL of liquid Ashdown medium and followed the enrichment protocol described above, while the other half was stored at −80°C for PCR analysis.

### DNA extraction and PCR analysis

The swab heads were cut and placed into the lysis buffer. The half filters, used for filtering the water samples, were ribolysed for three cycles of 20 s at 5,500 rpm (Lysing Matrix E two mL, MPBio, Eschwege, Germany) and DNA extraction was performed on 800 µL of the supernatant. Soil supernatants without pre-enrichment were extracted using the DNeasy PowerSoil Pro kit (QIAGEN, Hilden, Germany), while supernatants from enriched soil/water samples and rectal swabs were extracted using the High Pure PCR Template Preparation kit (Roche, Mannheim, Germany) according to the manufacturer’s instructions. To ensure extraction efficiency, an internal inhibition extraction control (Xeno™ DNA control or Diagenode IPC, Thermofisher) was added to each sample.

An initial PCR screening was performed using a real-time PCR assay targeting the *B. pseudomallei* complex and targeting *aro*A [30]. DNA samples that tested positive were further tested using real-time PCR assays targeting specific regions of the *B. pseudomallei* genome (*orf*11, BPSS0087, BPSS0745, and/or BPTT4176-4290) [31,32] and/or *B. thailandensis* (70 kDa) [33]. Each PCR reaction contained 5 µL of DNA and 15 µL of reaction mix, 1X of Universal Mastermix TaqMan^TM^ Fast Advanced 2X (Applied Biosystems^TM^, Vilnius, Lithuania), 0.5 µM of each primer, 0.1 µM of probe, 1X of IPC Diagenode and 2.8 µL of sterile water. The amplification procedure included a 2-min incubation at 50°C followed by a denaturation step at 95°C for 20 sec, then 45 cycles at 95°C for 3 sec and 60°C for 30 sec.

### Culture and screening of suspected colonies

Enrichment broths stored at 4°C were homogenized and 100 µL were plated on both conventional Ashdown agar and modified *B. cepacia* CHROMagar^TM^ agar (CHROMagar, Paris, France) supplemented with 4% glycerol, 500 mg/mL gentamicin and 130 mg/mL fosfomycin (CHR_GGF). Plates were incubated at 37°C and monitored daily. Suspect colonies (purple on Ashdown, green on CHR_GGF) were transferred to non-selective blood agar plates containing 5% horse serum (BA) and incubated at 37°C for 24 h. The bacterial suspensions were resuspended in sterile water, heated at 100°C for 20 min, and then centrifuged at 13,000 g for 10 min. The supernatant was used for PCR analysis (*aro*A and *orf*11 systems) to confirm colonies identification. Isolates confirmed to be *B. pseudomallei* by PCR were further characterized and their antimicrobial resistance pattern identified.

### B. pseudomallei strain characterization

#### Antibiogram

Antibiotic susceptibility testing was performed using the disk diffusion method according to European Committee on Antimicrobial Susceptibility Testing (EUCAST) guidelines [34]. The antibiotics tested included: amoxicillin-clavulanate (20-10 µg), meropenem (10 µg), trimethoprim-sulfamethoxazole (1.25−23.75 µg), chloramphenicol (30 µg), imipenem (10 µg), ceftazidime (10 µg), and tetracycline (30 µg) (Bio-Rad, France). Fresh 0.5 McFarland suspensions were prepared and plated on Mueller-Hinton agar (Bio-Rad, France). Antibiotic discs were then placed on the agar and plates were incubated at 35°C for 18 h. EUCAST breakpoints were used for interpretation. Two reference strains, *Escherichia coli* ATCC 25922 and *Staphylococcus aureus* ATCC 29213, were included as controls.

#### Biochemical testing

The API 20 NE system (Biomérieux, France) was used according to the manufacturer’s instructions. Bacterial suspensions were adjusted to a 0.5 McFarland standard and the API cassette was then incubated at 29°C for 24 h. The internet-based APIWEB database (accessed 7 April 2023, Biomérieux, France, https://apiweb.biomerieux.com/) was used to interpret the obtained profiles. Simultaneously, an oxidase test (oxidase reagent 50 x 0.75 mL, Biomérieux, France) was performed on each colony.

#### Multilocus sequence typing (*MLST)*

MLST was performed as previously described [35]. Amplified fragments were sent to Eurofins in Germany for sequencing. The sequences were compared to the *Burkholderia pseudomallei* PubMLST database for MLST profiling (https://pubmlst.org/https://pubmlst.org/organisms/burkholderia-pseudomallei, accessed 17 February 2023) [36].

## Results

### Serological survey in French Guiana and Guadeloupe

In French Guiana, seropositivity rates were 16% (58/361) in cattle, 2% (2/100) in goats, 3% (4/131) in sheep, 24% (15/63) in equids and 0% (0/15) in pigs (Table 1). The highest S/P% were observed in cattle, sheep and equids. In Les Saintes, 39% (12/31) of goats were seropositive (Table 1) and distributed over two farms (5/14 in farm A; 7/17 in farm B (Table 2)), with S/P% ranging from 0 to 503%.

**Table 1.** Summary of the serology (GLANDA-ELISA) results in livestock in French Guiana (n = 670) and Les Saintes, Guadeloupe (n = 31) S/P% serological titer expressed as the Sample to Positive ratio (calculated using a formula, see § Materials and Methods). Serum samples with S/P% ≥70% were considered positive. Farms where at least one animal tested GLANDA-ELISA positive were considered positive.

| Sampling site | French Guiana |  |  |  |  | Guadeloupe |
| --- | --- | --- | --- | --- | --- | --- |
| Animal species | Cattle | Goats | Sheep | Equids | Pigs | Goats |
| Number of positive animals/total (%) | 58/361 (16) | 2/100 (2) | 4/131 (3) | 15/63 (24) | 0/15 (0) | 12/31 (39) |
| [Min - Max] S/P% | [0-507] | [0-99] | [0-413] | [0-435] | [0-22] | [0-503] |
| Number of positive farms/total (%)* | 17/37 (45) | 2/10 (20) | 2/15 (13) | 4/6 (66) | 0/1 (0) | 2/2 (100) |

\* The number of farms in the table is higher than the total number of farms tested, as 10 farms included more than one animal species, i.e., appear in several columns.

**Table 2.**
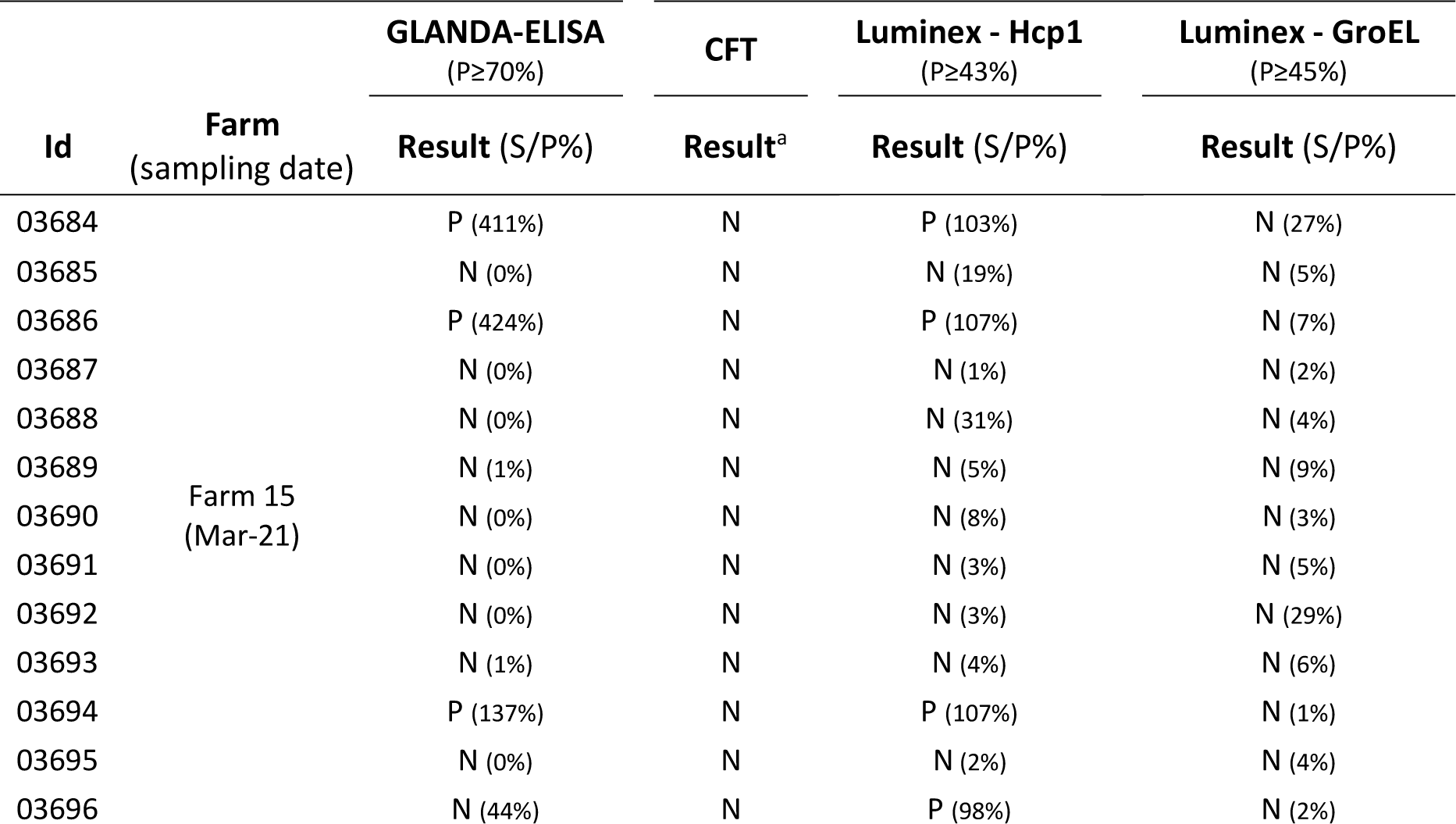
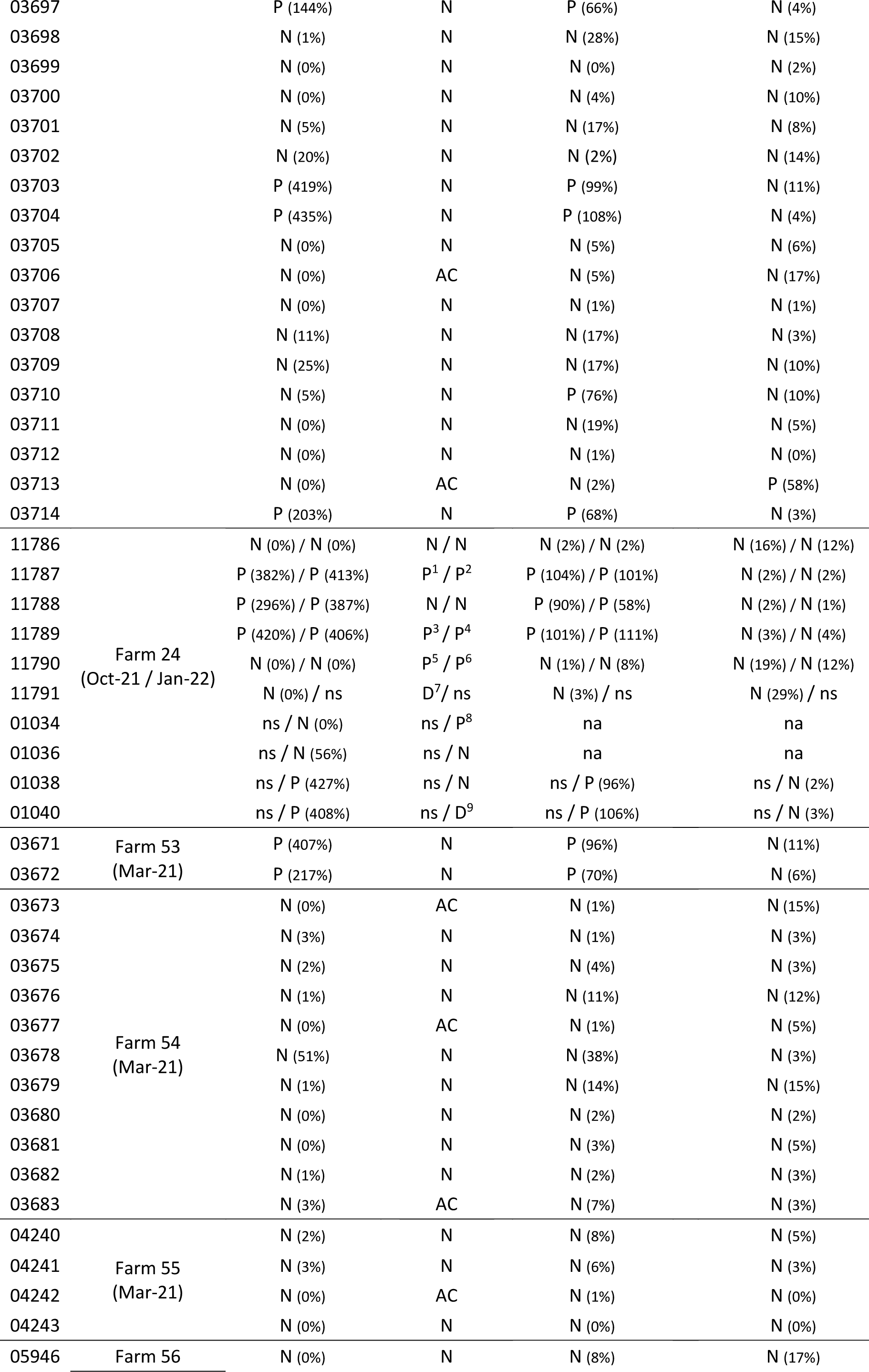
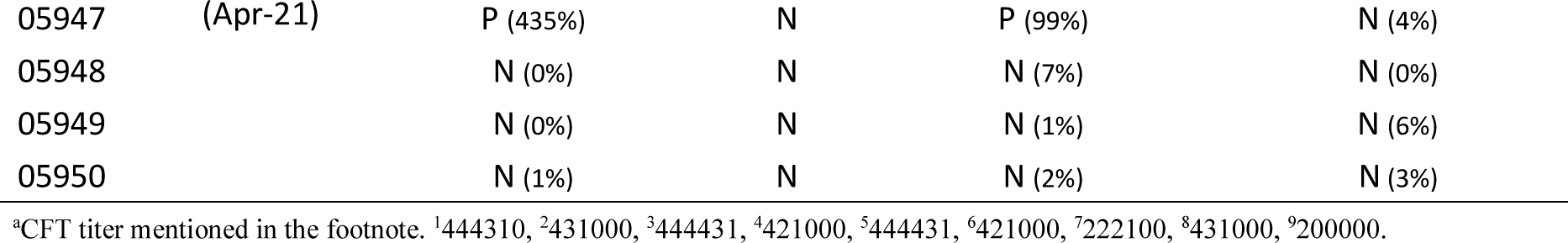
Serology results from 63 equids from six farms in French Guiana. GLANDA-ELISA: enzyme-linked immunosorbent assay with the ID Screen® Glanders Double Antigen Multi-species ELISA test (Innovative Diagnostics, Grabels, France). CFT: complement fixation test (see § Materiel and methods; Additional serological tests on equine sera, French Guiana). S/P%: serological titer expressed as the Sample to Positive ratio. Luminex: bead-based assay targeting recombinant proteins Hcp1 or GroEL. Id: animal identifier; P: positive; D: doubtful; N: negative; AC: anti-complementarity (interpretation not possible); na: not analysed; ns: not sampled.

In French Guiana, 25 out of 56 farms were tested positive, of which 10 included multiple species. Sixteen were in three townships: Macouria (n = 8), Kourou (n = 4), Sinnamary (n = 4). The serological results showed heterogeneity both between farms, differentiating positive and negative farms, as well as within positive farms, where both positive and negative animals were detected. Positive animals showed variable S/P% (**Fig 3, S1 Table**). For example, among the 25 ELISA-positive farms in French Guiana, 10 farms included animals with S/P% > 200.

**Fig 3.**
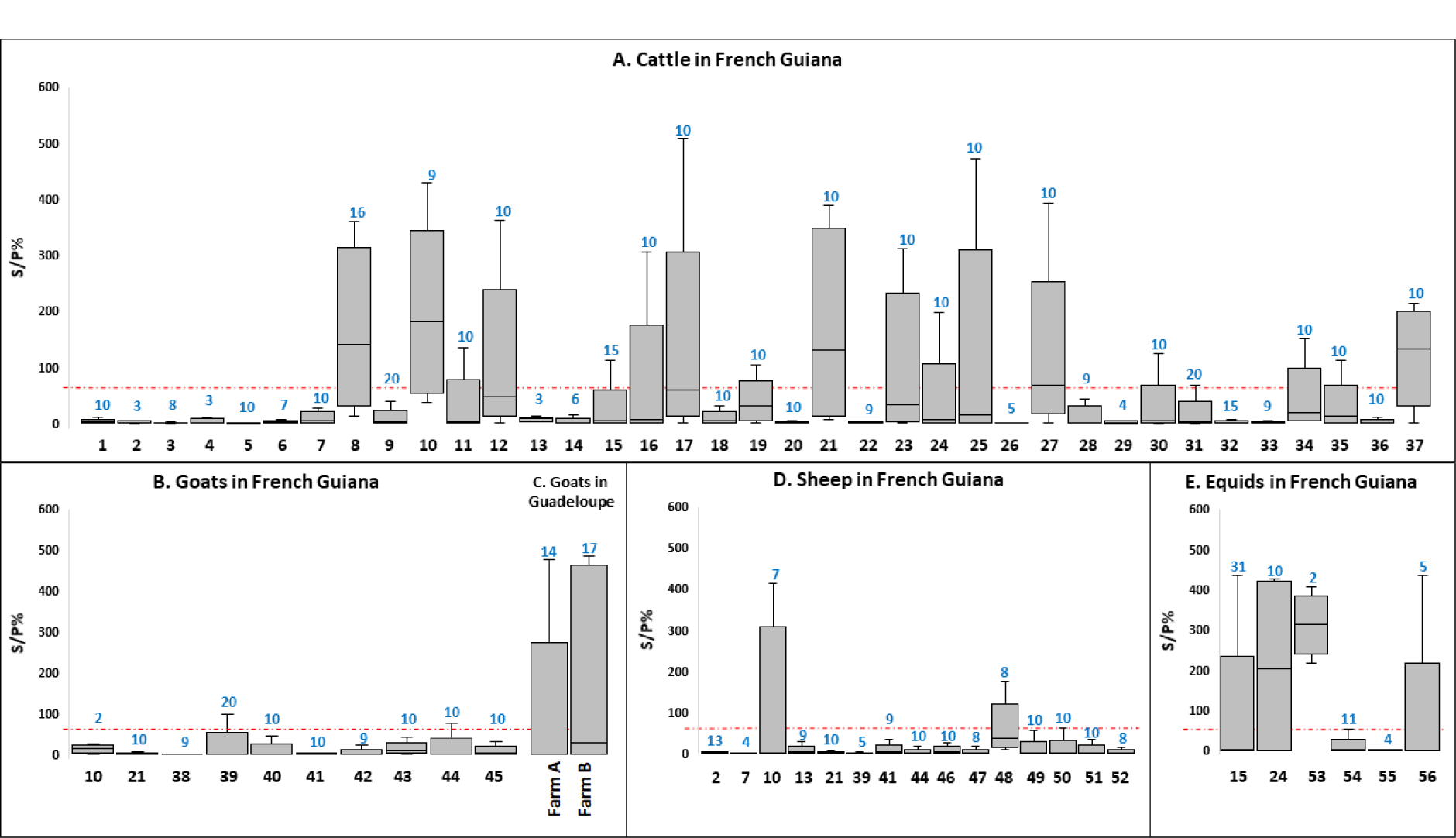
Summary of GLANDA-ELISA results per farm per animal species. (A) cattle (n = 37), (B) goats in French Guiana (n = 10), (C) goats in Les Saintes (n = 2), (D) sheep (n = 15), (E) equids (n = 6). S/P%: serological titer expressed as the Sample to Positive ratio. The x-axis shows the farm identification numbers. Farms are numbered from 1 to 56, when several species were on the same farm, the same number is used. The total number of animals tested per farm is shown in blue above each box plot. The red dotted line represents the S/P% threshold (70%). The bar above and below the box plot indicates the minimum and maximum S/P%. The top and bottom of the box indicate the 1^st^ and 3^rd^ quartiles of S/P%. The bar in the box indicates the median. Due to the small number of pig farms tested, we have excluded these data from the following figure.

### Additional analysis on equids from French Guiana and follow-up on farm #24

In French Guiana, equids originating from six farms out of 56 (2 to 31 equids per farm; n=63) were further tested by two additional methods developed for the diagnosis of glanders: the reference CFT method and the newly developed Luminex method using GroEL and Hcp1 antigens [28]. All equids with ELISA-positive results were confirmed positive with Hcp1 by Luminex, but negative with CFT and with GroEL by Luminex except for farm 24 (**Table 2**). In this farm, some of the horses/equids were tested positive with CFT.

### Focused study in two goat farms in Les Saintes, Guadeloupe

#### Serological and molecular biology study

To investigate the temporal evolution of the serological response in two goat farms that initially tested ELISA-positive in November 2021, additional blood samples were collected in April 2022, August 2022, and December 2022. Farm A was also sampled again in March 2023.

Seropositive animals were consistently detected in both farms at all-time points, although the percentage of positive animals varied over time. It ranged from 13% to 36% in farm A and from 23% to 41% in farm B. The highest number of ELISA-positive animals was observed in both farms in November 2021 (Table 3).

**Table 3.**
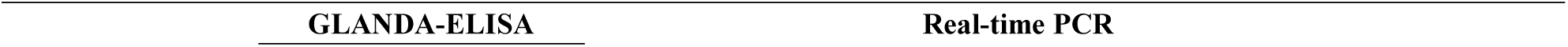
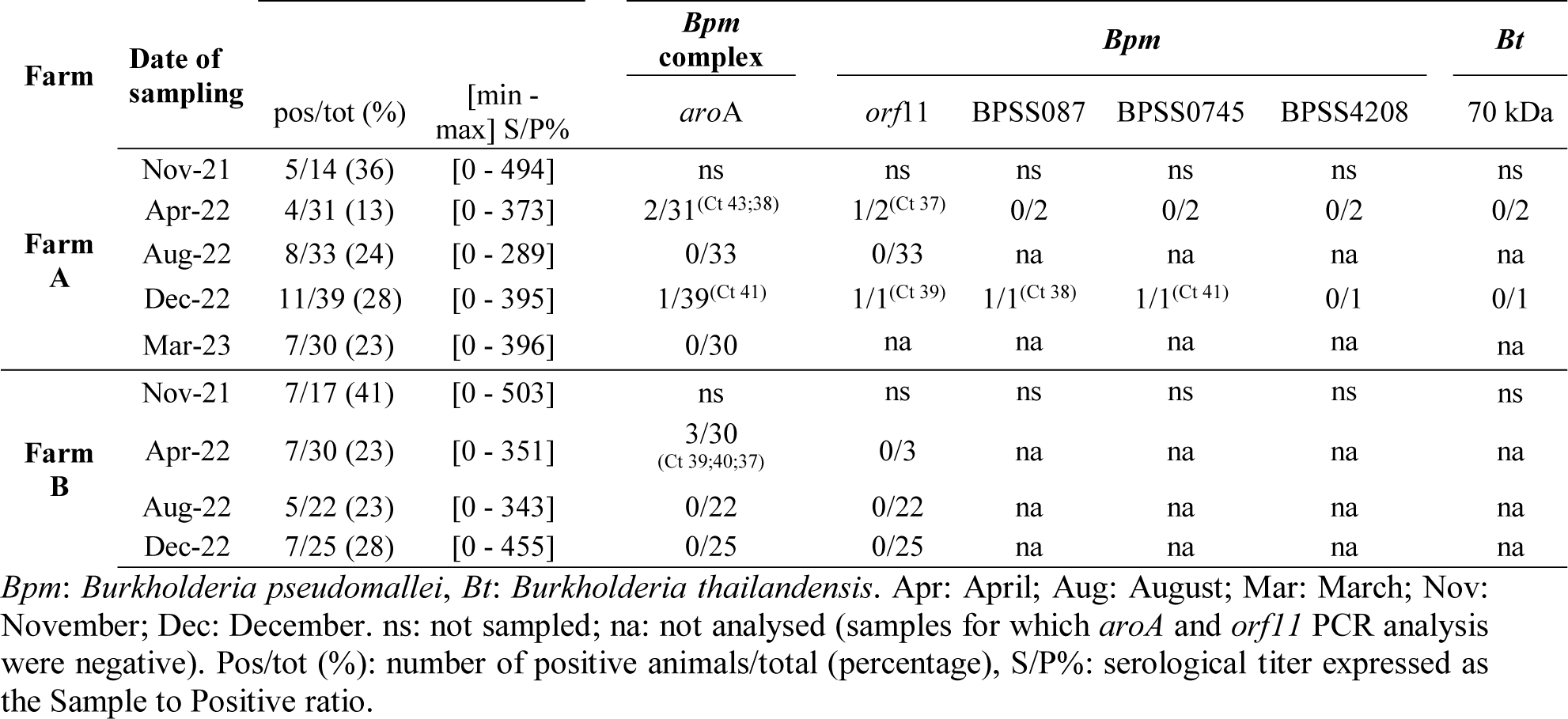
Serological and molecular results from two goat farms in Les Saintes, Guadeloupe. The ratio of real-time PCR-positive rectal swabs samples to the total number of samples tested is provided for six gene targets. The PCR cycle threshold (Ct) is provided for positive samples. ELISA: enzyme-linked immunosorbent assay with the ID Screen® Glanders Double Antigen Multi-species ELISA test (Innovative Diagnostics, Grabels, France).

| Farm | Date of sampling |  |  | *Bpm* complex | *Bpm* |  |  |  | *Bt* |
| --- | --- | --- | --- | --- | --- | --- | --- | --- | --- |
|  |  | pos/tot (%) | [min - max] S/P% | *aroA* | *orf11* | BPSS087 | BPSS0745 | BPSS4208 | 70 kDa |
| Farm A | Nov-21 | 5/14 (36) | [0 - 494] | ns | ns | ns | ns | ns | ns |
|  | Apr-22 | 4/31 (13) | [0 - 373] | 2/31<sup>(Ct 43;38)</sup> | 1/2<sup>(Ct 37)</sup> | 0/2 | 0/2 | 0/2 | 0/2 |
|  | Aug-22 | 8/33 (24) | [0 - 289] | 0/33 | 0/33 | na | na | na | na |
|  | Dec-22 | 11/39 (28) | [0 - 395] | 1/39<sup>(Ct 41)</sup> | 1/1<sup>(Ct 39)</sup> | 1/1<sup>(Ct 38)</sup> | 1/1<sup>(Ct 41)</sup> | 0/1 | 0/1 |
|  | Mar-23 | 7/30 (23) | [0 - 396] | 0/30 | na | na | na | na | na |
| Farm B | Nov-21 | 7/17 (41) | [0 - 503] | ns | ns | ns | ns | ns | ns |
|  | Apr-22 | 7/30 (23) | [0 - 351] | 3/30<sup>(Ct 39;40;37)</sup> | 0/3 | na | na | na | na |
|  | Aug-22 | 5/22 (23) | [0 - 343] | 0/22 | 0/22 | na | na | na | na |
|  | Dec-22 | 7/25 (28) | [0 - 455] | 0/25 | 0/25 | na | na | na | na |

*Bpm*: *Burkholderia pseudomallei*, *Bt*: *Burkholderia thailandensis*. Apr: April; Aug: August; Mar: March; Nov: November; Dec: December. ns: not sampled; na: not analysed (samples for which *aroA* and *orf11* PCR analysis were negative). Pos/tot (%): number of positive animals/total (percentage), S/P%: serological titer expressed as the Sample to Positive ratio.

Rectal swabs were collected in parallel with blood samples to test for the fecal shedding of *B. pseudomallei*. In total, six swabs tested positive by PCR for the *B. pseudomallei* complex, of which two were positive for *B. pseudomallei* species, one collected in April 2022 and one in December 2022 in farm A (Table 3).

The longitudinal serological study started in August 2022, when as many goats as possible were implanted with a microchip to identify individual goats. In farm A, 20 goats were sampled three times: 13 were consistently ELISA-negative, five were consistently ELISA-positive and two changed status during the course of the 8-month study (**Table 4**). Of note, six goats had S/P% greater than 200% on at least two sampling events. One goat with the lowest S/P% (≈150%) later tested negative in March 2023. Conversely, two goats with initially low S/P% later had higher values. Of the seven goats that were seropositive at some point, five were over one year of age. The two identified mother-daughter pairs were consistently negative.

**Table 4.**
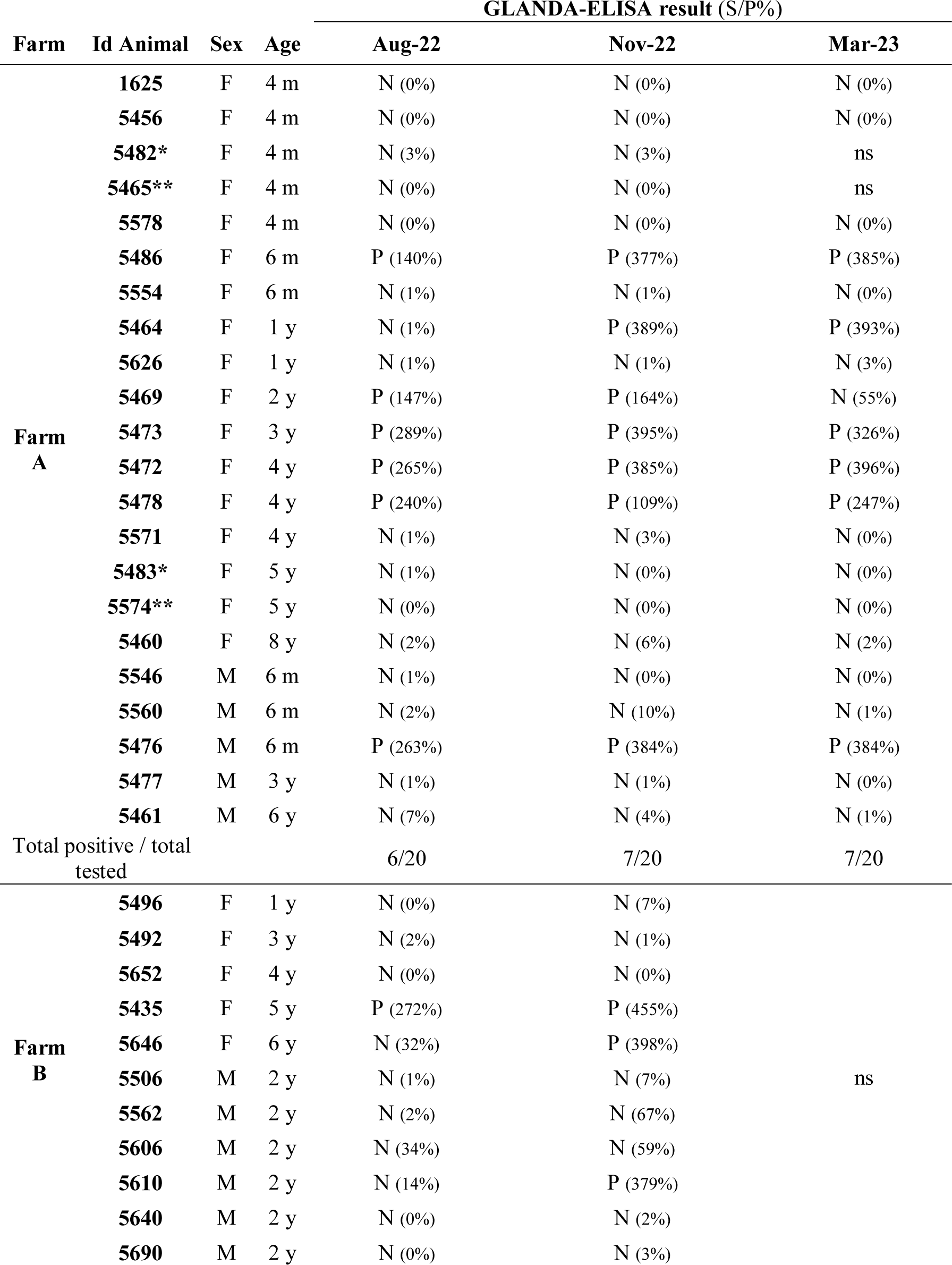
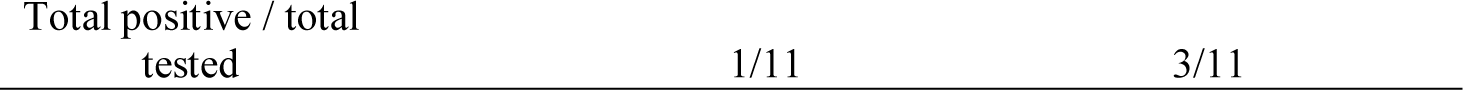
Longitudinal serological study in two goat farms in Les Saintes, Guadeloupe. GLANDA-ELISA: enzyme-linked immunosorbent assay with the ID Screen® Glanders Double Antigen Multi-species ELISA test (Innovative Diagnostics, Grabels, France). S/P%: serological titer expressed as the Sample to Positive ratio, N: negative (S/P% <70), P: positive (S/P% ≥70%), F: female, M: male, m: month old, y: year-old, ns: not sampled (farm B was not sampled in March 2023). * 5483 is the mother of 5482; ** 5574 is the mother of 5465.

In farm B, 11 animals were sampled in August and December 2022. Eight tested ELISA-negative twice, one tested ELISA-positive twice, and two changed status during the course of the study (seroconversion). All three animals that were seropositive at some points were over one year of age and had S/P% values greater than 200%.

#### Environmental sampling in goat pastures

In December 2022, 50 soil samples and two water samples (from the drinking troughs) were collected from farm A; and 42 soil samples and eight water samples (from the drinking troughs) were collected from farm B. All samples were analysed by PCR and bacteriological culture. Samples from farm A were all PCR-negative, pre- or post-enrichment. In farm B, two out of 42 soil samples (#313 and #336) were PCR-positive for the *B. pseudomallei* complex post-enrichment, but only sample #313 tested PCR-positive for *B. pseudomallei* (all four specific targets) (**Table 5**).

**Table 5.** Results from molecular testing of environmental samples from two goat farms in Les Saintes, Guadeloupe. Soil and water samples collected in December 2022 were tested by real-time PCR for the presence of different *Burkholderia* spp. Results are provided as the ratio of positive samples against the total number of tested samples tested for each target. For PCR-positive results, Ct values are provided. na: not analysed.

| Farm | Type of sample | Number of samples | *Bpm complex* | *Bpm* |  |  |  | *Bt* |
| --- | --- | --- | --- | --- | --- | --- | --- | --- |
|  |  |  | *Bpm*<br>aroA | orf11 | BPSS087 | BPSS0745 | BPSS4208 | 70 kDa |
| Farm A | soil | 50 | 0/50* | na | na | na | na | na |
|  | water | 2 | 0/2 | na | na | na | na | na |
| Farm B | soil | 42 | 2/42* (Ct 34 ;34) | 1/2*(Ct 40) | 1/2*(Ct 38) | 1/2*(Ct 37) | 1/2*(Ct 39) | 0/2* |
|  | water | 8 | 0/8 | na | na | na | na | na |

*Bpm*: *Burkholderia pseudomallei*, *Bt*: *Burkholderia thailandensis*. \* PCR results were obtained post-enrichment (see § Material and Methods).

PCR-positive soil samples #313 and #336 were plated post-enrichment on both Ashdown and CHR_GGF agar media for isolation. A total of 46 suspect colonies were isolated from soil #313. Of these, 30 were PCR-positive for the *B. pseudomallei* complex and all four *B. pseudomallei* specific targets. Fifteen suspect colonies were isolated from soil #336 and tested PCR-positive for the *B. pseudomallei* complex but PCR-negative for *B. pseudomallei* and *B. thailandensis*.

*B. pseudomallei* colonies on Ashdown initially appeared purple and smooth after 24 h at 37°C and then changed to a rough, dry texture after 48 h (**Fig 4A**). On CHR_GGF, the same morphological changes were observed: colonies were green and smooth after 24 h of culture, but changed to a white, rough, and dry texture after 120 h (**Fig 4B**).

**Fig 4.**
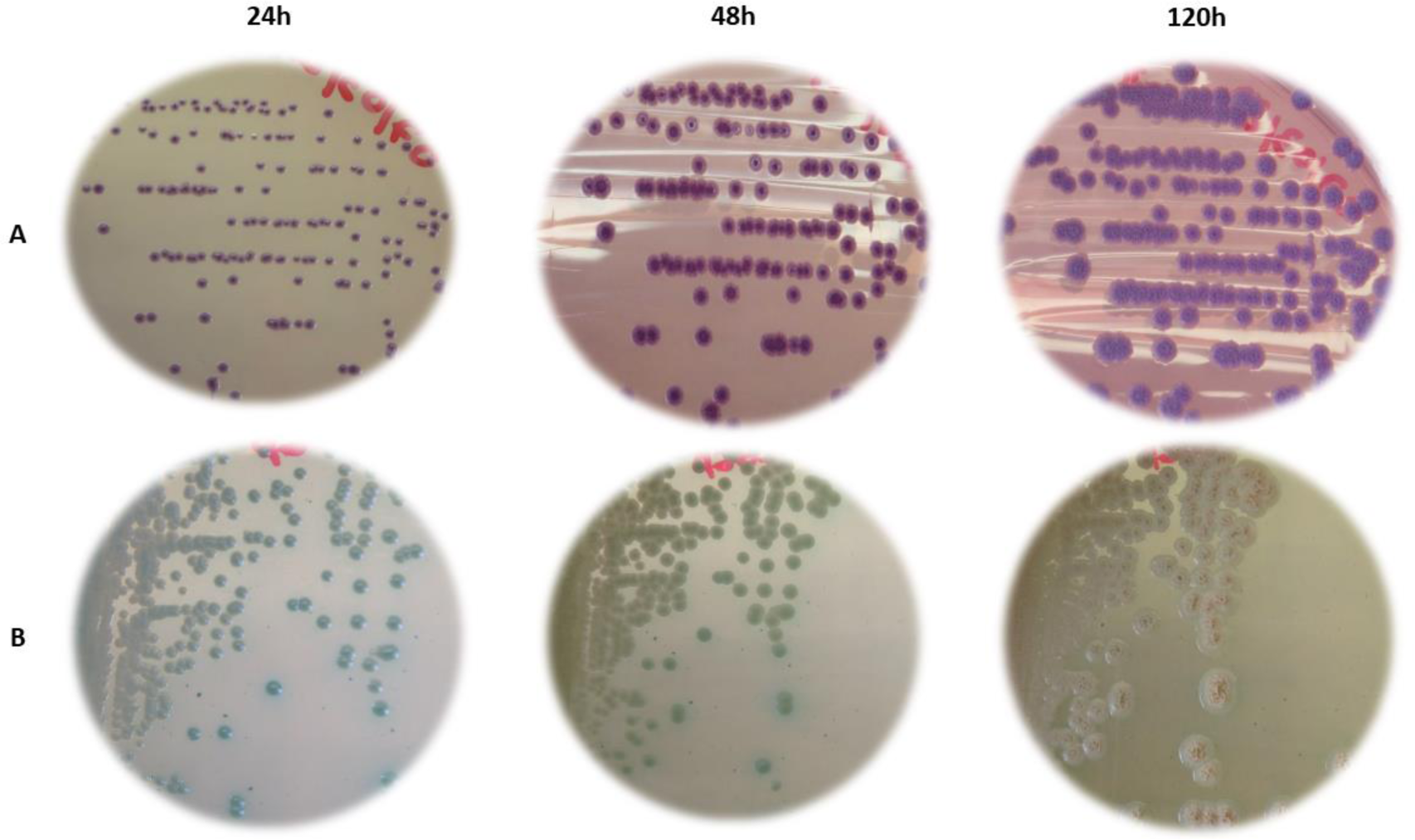
*Burkholderia pseudomallei* strain D22-10884_313#20 cultures. Photographs from culture plates (A) on Ashdown and (B) on CHR_GGF (modified *B. cepacia* CHROMagar^TM^ agar supplemented with 4% glycerol, 500 mg/mL gentamicin and 130 mg/mL fosfomycin) after a 24, 48 and 120 h incubation at 37°C. The strain was isolated from soil #313 collected from farm B in Les Saintes, Guadeloupe.

One *B. pseudomallei* colony (D22-10884_S313#20) isolated from soil #313 was selected for further analysis. The strain showed the API 20 NE profile 1-456-574 together with a positive result for the oxidase test. When compared with the profiles listed in APIWEB, the strain had the highest identification scores with *B. pseudomallei* (47.9%) and *B. cepacia* (45.3%). The antibiotic susceptibility profile of strain D22-10884_S313#20 showed sensitivity to all antibiotics tested (commonly used for human melioidosis treatment): amoxicillin-clavulanate, ceftazidime, tetracycline, imipenem, meropenem, trimethoprim-sulfamethoxazole and chloramphenicol (**S2 Table**). The MLST sequence type of strain D22-10884_S313#20 was ST92.

## Discussion

By interpreting positive ELISA results in animals as a sign of previous environmental exposure to *B. pseudomallei*, our serological screening showed variable results between regions but also between species. In French Guiana, the seroprevalence was 2% in sheep and goats, 15% in cattle and 22% in equids; in Les Saintes, where only goats were tested, the seroprevalence was 39%. In comparison, studies carried out in endemic countries using different serological methods showed seroprevalence of 6% and 13.6% in sheep, 0.3% and 2.6% in goats, and 2% and 7.6% in cattle, in Thailand (using IHA) [37] and Malaysia (using CFT) [38], respectively. Due to different levels of endemicity and methodological strategies, the data are not easily comparable. Our serological screening also showed within-farm variability. The longitudinal study conducted in two ELISA-positive farms in Les Saintes showed that most goats remained ELISA-positive over an 8-month period. Further studies are needed to better understand the long-term immune response dynamics after exposure to *B. pseudomallei*. This should help to explain the within-farm heterogeneity, i.e., why animals living in farms with similar environmental exposure do not all show the same serological result.

Melioidosis has been studied primarily in well-established endemic regions such as Southeast Asia and Australia with limited research in Central and South America [26,39]. In the Caribbean, human cases have been documented for over 30 years in Guadeloupe and Martinique [21–23,25] but the local presence of *B. pseudomallei* in the environment has never been established. Isolating the strain from the environment is challenging, as demonstrated by a recent environmental survey in Puerto Rico where only three out of 500 soil samples from 60 sites tested positive for *B. pseudomallei* using PCR, with only one sample being culture positive [40]. Previous exposure to *B. pseudomallei* promotes an immune response in animals that can be detected serologically. Hence, one major objective of our study was to confirm the benefit of an initial serological screening in animals to increase the probability of detecting *B. pseudomallei* in the environment.

The GLANDA-ELISA kit, originally developed for the diagnosis of equine glanders [11], was used on a goat farm in New Caledonia where one case of melioidosis was confirmed. The serological survey allowed to confirm exposure in 35% of goats on this farm, while those from three other farms with no clinical case were 100% seronegative (Laroucau *et al.,* en révision). However, the validation of this ELISA kit in detecting clinical melioidosis and/or previous exposure to *B. pseudomallei* called for further data. The application of the GLANDA-ELISA kit in our multi-species study yielded heterogeneous results between individuals and between farms, with some farms fully seronegative. Overall, serology results were consistent with the absence (in French Guiana) or limited number (in Les Saintes) of human melioidosis cases.

Our study supports the idea that animals could be used as sentinels of *B. pseudomallei* presence in the environment and that the GLANDA-ELISA kit is a relevant tool to do so. Further research is now necessary to establish the specificity of this kit in a context different from its original design and validation. The high diversity and prevalence of *Burkholderia* species in the environment, including closely related species within the *B. pseudomallei* complex, raises concerns about the possibility of cross-reactivity. Since the identity of the protein used in the GLANDA-ELISA kit is not disclosed, it is not possible to assess its degree of conservation in environmental *Burkholderia* strains, if any, and thus the risk of cross-reactivity when using this kit. In addition, the cutoff used in our study was originally established for equine glanders in a clinical context and not for the assessment of environmental exposure to *B. pseudomallei*. A reevaluation of the cutoff for our specific goal and in different host species may help to refine the data obtained with this kit.

While quantitative data on *B. pseudomallei* in the environment are scarce, it is plausible that in non-endemic areas the bacteria are present at lower concentrations which might decrease the probability of detection. In anticipation of this additional challenge, we conducted our environmental sampling in Les Saintes during the rainy season (June to November), a period identified as the most favorable for the presence of *B. pseudomallei* in the soil [15]. We also optimized the protocol from existing guidelines [18], adding two new media to the conventionally recommended Ashdown medium [29]. First, an erythritol-based medium was used as the sole carbon source during the second step of a two-step enrichment to select for *B. pseudomallei* [19]. A similar two-step enrichment protocol (TBSS-C50 + erythritol) recently allowed the isolation of environmental *B. pseudomallei* strains in Ghana, West Africa [20]. Second, we used a chromogenic medium originally developed for the detection of *B. cepacia* in clinical samples as green colonies. To increase its selectivity for *B. pseudomallei*, we modified the medium by incorporating new antibiotics and confirmed that reference strains of *B. pseudomallei* grew well on this medium (S1 Figure). To our knowledge, this is the first use of a chromogenic medium to improve the selection of *B. pseudomallei* colonies from poly-contaminated environmental matrices. Despite these protocol improvements, in our study only one soil sample (out of 92 samples collected from seropositive farms) and a single *B. pseudomallei* strain was isolated. Although the use of media CHROMagar^TM^ *B. cepacia* seems promising, further protocol optimization (e.g., sample volume, enrichment time, chromogenic and/or selective medium for environmental matrices) will be needed to facilitate the detection and isolation of environmental strains of *B. pseudomallei.* Of note, an additional soil sample tested PCR-positive for the *B. pseudomallei* complex, but not for *B. pseudomallei* or *B. thailandensis* species. Further characterization of all non-*B. pseudomallei* strains isolated in this study will help identify all the *Burkholderia* species to which animals can be exposed and to assess their cross-reactivity with the GLANDA-ELISA kit.

Differentiating glanders (caused by *B. mallei*) from melioidosis (caused by *B. pseudomallei*) is challenging due to their high antigenic similarity [13], with *B. mallei* being a monophyletic clade within *B. pseudomallei* [41]. Considering that (i) the GLANDA-ELISA kit has been designed for the diagnosis of equine glanders, and that (ii) glanders is frequently reported in Brazil [42], a country sharing a border with French Guiana, the detection of GLANDA-ELISA positive horses in French Guiana warranted further testing. Clinically healthy horses that tested ELISA-positive were all Hcp1-positive by Luminex but GroEL-negative by Luminex, both proteins being key markers for glanders diagnosis [28]. When adding CFT (the reference method for glanders diagnosis) as a fourth test, all ELISA-positive horses were CFT-negative, except for horses on farm #24. In this farm, the detection of horses positive for three out of four tests relevant to glanders diagnosis prompted close surveillance. A follow-up examination three months later confirmed the initial serological results as well as the absence of clinical signs typical of acute glanders (e.g., respiratory distress or abscesses). Overall, the variability in test results among animals in farm #24 may indicate different exposure and/or infection dynamics and/or test characteristics. Despite the absence of previously reported cases of glanders or melioidosis in French Guiana, the final health status of these equids remains uncertain given the chronic potential of these two diseases [43–45].

Although the serological approach is generally not favored for the epidemiology of melioidosis in endemic areas, due to more frequent environmental exposure of humans and animals and the resulting challenges of interpreting positive results outside of a clinical context [46,47], this approach can be promising for non-endemic areas. The GLANDA-ELISA kit may be particularly valuable for identifying past exposure in animals and conducting targeted environmental surveys around exposed animals. This is particularly relevant for livestock that are often confined to restricted areas.

Our serological and molecular screening on two goat farms in Les Saintes, followed by an environmental survey, confirmed fecal shedding of *B. pseudomallei* in one farm and successfully led to the isolation of an environmental strain of *B. pseudomallei* in the other farm. These results provide the first definite confirmation of the local establishment of *B. pseudomallei* in Les Saintes, Guadeloupe. They also support the hypothesis of a local contamination of two cases of human melioidosis reported on the island in 1997 (a tourist that visited Les Saintes for 3 weeks) [25] and in 2016 (a local resident) [23]. Importantly, the MLST profile of the strain isolated from our study (ST92) was identical to that of the 1997 human case [23]. In the *Burkholderia pseudomallei* PubMLST database, ST92 was identified in nine human strains: one from a Swiss traveler to Martinique [21], one from Puerto Rico, two from Mexico [48], and five from Brazil [49], illustrating the restricted spatial distribution of this sequence type.

Even though human melioidosis has not been reported so far in French Guiana, our results call for the implementation of environmental surveys around the ELISA-positive farms identified in this study, as well as a retrospective re-evaluation of human infections with melioidosis-like symptoms. In Guadeloupe and Martinique, where several human cases have been diagnosed in recent years [23,24], we suggest adding melioidosis to existing animal surveillance programs.

## Acknowledgements

This study was supported by the Ile de France Region (Dim1Health) and the European Commission Directorate-General for Health and Consumers (EC no.180/2008). We thank the French Institut de Recherche pour le Développement (IRD) and the French Agence nationale de sécurité sanitaire de l’alimentation, de l’environnement et du travail “ANSES” for their financial support. We also thank all the livestock farmers, particularly the owners of the two goat farms in Les Saintes, who allowed repeated sampling to complete the longitudinal study. Special thanks to CHROMagar^TM^ who supplied the culture media and Dr. Sophie Granier for her valuable advice on the implementation of the antibiogram test. This work is part of the Doctoral requirements of the first author (MG).

## Conflicts of Interest

None of the authors have any financial or personal relationships with any individuals or organizations that could inappropriately influence or bias this paper.

## Author Contributions

Conceptualization: Karine Laroucau (KL), Emma Rochelle-Newall (ERN), Vanina Guernier-Cambert (VGC), Sébastien Breurec (SB)

Resources: Gil Manuel (GM), Xavier Baudrimont (XB), Sébastien Breurec (SB) Methodology: Mégane Gasqué (MG), Karine Laroucau (KL)

Investigation: Mégane Gasqué (MG), Gil Manuel (GM), Xavier Baudrimont (XB), Rachid Aaziz (RA), Thomas Deshayes (TD), Jules Terret (JT)

Supervision: Karine Laroucau (KL), Emma Rochelle-Newall (ERN), Vanina Guernier-Cambert (VGC)

Writing –original draft: Mégane Gasqué (MG), Karine Laroucau (KL)

Writing –review & editing: Mégane Gasqué (MG), Karine Laroucau (KL), Emma Rochelle-Newall (ERN), Vanina Guernier-Cambert (VGC), Gil Manuel (GM), Xavier Baudrimont (XB), Rachid Aaziz (RA), Thomas Deshayes (TD), Jules Terret (JT), Sébastien Breurec (SB)

## Supporting information

**S1 Table.** Serological results of animals in French Guiana and Les Saintes (Guadeloupe)

**S2 Table.** Antibiogram results of 22-10884_313#20 strains

**S1 Figure**. Evaluation of CHROMagar^TM^ *B. cepacia* as a relevant media for other *Burkholderia* strains.

## Notes

### Competing Interest Statement

The authors have declared no competing interest.

