## Supplementary figures and images for "Serological screening in animals combined with environmental surveys provides definite proof of the local establishment of *Burkholderia pseudomallei* in Guadeloupe"

### Supplemental Figure 1

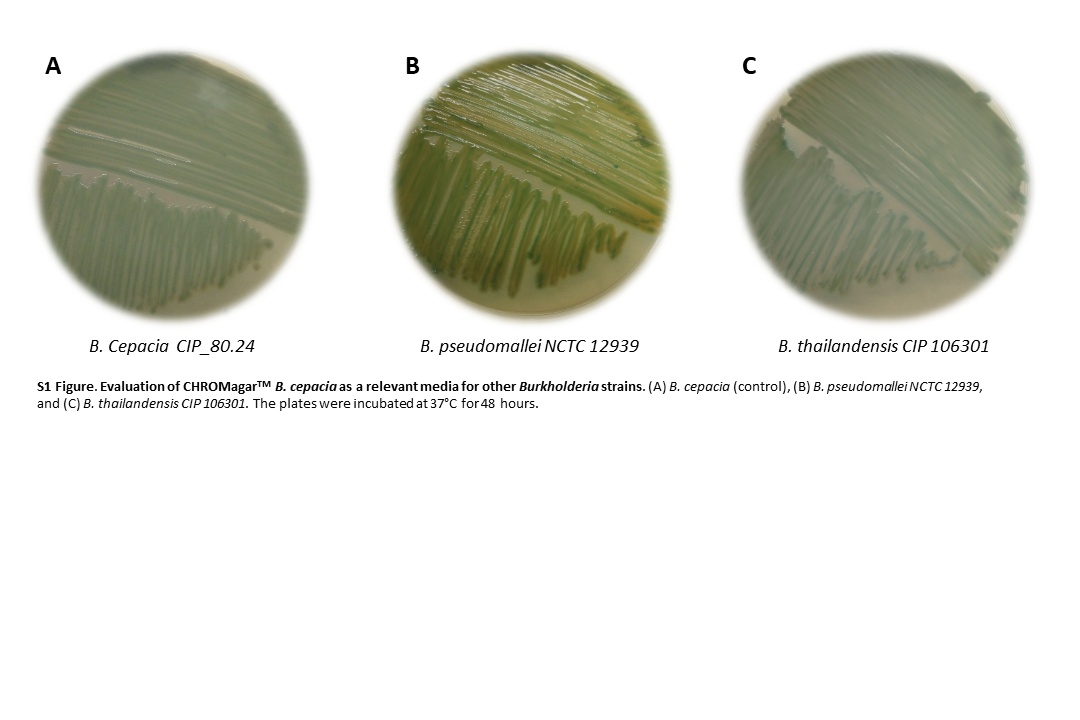
